# Amyloid-beta is present in the spinal cord of APP/PS1 mice and may contribute to neuropathology manifesting as lower urinary tract dysfunction

**DOI:** 10.64898/2026.07.09.737006

**Authors:** Sophia M. Vrba, Ajinkya R. Limkar, Kimberly Keil-Steitz, Jeffrey J. Nirschl, Collin J. Laaker, Dhruv Bansal, Simon Federico Woen Ordonez, Erin G. Brooks, Jeffrey Helgager, Mariana Pehar, Matyas Sandor, William A. Ricke, Zsuzsanna Fabry

## Abstract

Urinary incontinence (UI) is a common and debilitating comorbidity in Alzheimer’s disease (AD), yet its underlying pathophysiology remains poorly defined. While UI in dementia has traditionally been attributed to functional impairment, emerging clinical and urodynamic data suggest that neurologic mechanisms may contribute to lower urinary tract dysfunction in this population. Here, we investigated urinary function and neuropathological changes in aged APP/PS1 mice (AD mice), a widely used model of amyloid pathology. Using functional voiding assays, we identified a pattern of urinary dysfunction characterized by increased urinary frequency, small-volume voiding, shortened void duration, and reduced bladder compliance in the absence of bladder outlet obstruction or gross changes in bladder or prostate morphology. These findings are most consistent with a storage-phase abnormality accompanied by impaired voiding coordination rather than classic detrusor overactivity or underactivity. We examined spinal cord and peripheral components involved in bladder innervation and identified amyloid-beta deposition throughout the thoracolumbar and lumbosacral spinal cord, dorsal root ganglia, ventral roots, cauda equina, and associated meningeal structures in AD mice. Importantly, amyloid deposition was accompanied by reduced expression of vesicular acetylcholine transporter and decreased neuronal activation in bladder-innervating pathways, without evidence of increased apoptosis. Taken together, these data demonstrate that AD mice develop a mixed lower urinary tract dysfunction phenotype associated with amyloid-beta deposition and altered neuronal signaling within the spinal cord and peripheral micturition pathways. These findings support a neurogenic contribution to urinary dysfunction in AD and highlight the spinal cord as a novel site of pathology that may influence urinary symptoms in Alzheimer’s dementia.

## Introduction

Alzheimer’s disease (AD) is the most common form of dementia, affecting 65 million individuals worldwide, with the prevalence expected to increase to 82 million by 2050[1]. AD is a chronic, progressive neurodegenerative disorder characterized by extracellular deposition of amyloid-beta plaques and intracellular neurofibrillary tangles comprised of hyperphosphorylated tau protein. There has been robust research on the pathogenesis of AD within the brain parenchyma, but few studies have focused on pathological involvement of Alzheimer’s disease outside of the neuropathologic effects in the brain. The spinal cord is part of the central nervous system and is a key innervation pathway relaying sensory, motor, and reflex information from the periphery to the brain. However, neuropathological changes within the spinal cord have been far less studied in the context of AD, and there is evidence suggesting the involvement of spinal cord innervation pathways in the systemic manifestations of AD. MRI studies, only within the cervical spinal cord, have shown that AD patients have more atrophy of the spinal cord in comparison to healthy control subjects [2]. Early neuropathologic analysis has found that amyloid-beta was present within the spinal cord, but which section of the cord was not specified [3]. Although amyloid-beta plaques have been observed in the spinal cord in preclinical models of AD [4], the role and pathological significance of amyloid-beta accumulation in extra-cerebral regions such as the spinal cord remain understudied. Here, we described micturition dysfunction associated with amyloid-beta deposition within the lumbosacral sacral spinal cord of aged APP/PS1 mice, accompanied by evidence of altered neuronal state and dysregulated spinal innervation pathways.

Urinary incontinence (UI), defined as involuntary leakage of urine, is a common comorbidity in patients with AD, and individuals with dementia are at a significantly higher risk for developing UI even when controlling for age[5–8]. These studies are limited in that the studies did not control for the presence of being on acetylcholinesterase inhibitors (AChEIs) [5–8]. UI negatively affects health-related quality of life measures, with affected patients reporting increased social isolation and higher rates of depression[9, 10]. In a study of 75 patients with dementia and their families and caregivers, 83% of patients reported limitations in daily activities due to UI, and all families/or caregivers reported that UI influenced the care they provided[11]. Importantly, UI is recognized as a major risk factor for nursing home placement in this population[12].

The clinical symptoms and underlying drivers of UI in the context of neurodegeneration are complex and multifactorial. Historically, UI in AD was largely attributed to functional urinary incontinence, wherein patients retain bladder sensation but are unable to respond appropriately due to impaired cognition, reduced motivation, or reduced mobility[13, 14]. Functional UI is not caused by intrinsic pathology in the lower urinary tract but rather reflects disease progression in the context of cognition or behavior. However, a growing body of research suggests that UI in AD is heterogeneous and arises from a combination of functional deficits and pathological changes associated with the disease process. Urodynamic studies have suggested that urge incontinence, driven by detrusor overactivity, is the most prevalent form of UI in AD patients, present in 57.6% of patients[13, 15, 16]. These studies also identified mixed incontinence and stress incontinence in a subset of patients, emphasizing the importance of a comprehensive clinical evaluation to account for the diverse etiologies of UI in patients with dementia. Despite the increasing recognition that UI in dementia may be driven by neurodegenerative pathology, relatively few studies have directly examined the underlying mechanisms contributing to urinary dysfunction in AD.

To further complicate clinical management, UI and dementia pose opposing therapeutic challenges due to conflicting pharmacologic mechanisms of mainstream treatments. Early neurostructural changes in AD are characterized by degeneration of cholinergic nuclei in the basal forebrain and cerebral cortex[17]. As acetylcholine is an essential neural transmitter in memory and cognitive function, AChEIs, such as donepezil or rivastigmine, are commonly prescribed to symptomatically treat AD by preventing acetylcholine degradation and increasing synaptic availability[18, 19]. Similarly, the urinary bladder relies on cholinergic signaling during the voiding phase; therefore, increased synaptic acetylcholine has been shown to increase risk of overactive bladder by worsening lower urinary tract symptoms[20]. In contrast, anticholinergic medications, such as oxybutynin and tolterodine, represent the gold standard for UI management as they inhibit cholinergic signaling at the muscarinic receptors in the detrusor muscle to reduce involuntary contractions [21]. Notably, anticholinergic medication exposure has been associated with an increased risk of dementia in older adults[22]. Given the high prevalence of UI in AD and this therapeutic paradox, additional studies to elucidate the pathophysiology of UI in AD are essential to reveal new therapeutic targets and treatment avenues for these patients.

In this study, we investigated and characterized the contribution of spinal cord dysfunction in the development of urinary dysfunction in 24-month-old (aged) male APP/PS1 mice, a well-established model of amyloid-beta deposition. We demonstrated that aged male APP/PS1 mice exhibit increased lower urinary tract dysfunction when compared to their wild-type (WT) controls. The mice displayed shortened void durations with preserved flow rate, reduced bladder compliance, and no changes in bladder mass, volume, or prostate lobe enlargement. When examining spinal cords from aged male APP/PS1 mice, we found amyloid-beta deposition throughout the neuroaxis, including the cervical, thoracic, lumbar, and sacral spinal cord, as well as within lumbosacral structures involved in bladder sensory and autonomic signaling, including the dorsal root ganglia, conus medullaris, and cauda equina. Consistent with these findings, analyses of neuronal activity revealed reduced vesicular acetylcholine transporter (VAchT) and decreased c-Fos labeling in spinal cord regions involved in bladder, internal urethral sphincter, and external urethral sphincter control in aged APP/PS1, Collectively, these findings identify spinal cord amyloid-beta pathology and associated nerve dysfunction as a potential contributor to urinary dysfunction in the setting of AD.

## Methods and materials

### Animals

All animal experiments were conducted under the protocols approved by the University of Wisconsin Animal Care and Use Committee. 24-month-old male B6.Cg-Tg(APPswe,PSEN1dE9)85Dbo/Mmjax (APP/PS1) mice and their WT littler mate controls were obtained from the Jackson Laboratory. Mice were maintained as hemizygous animals in a congenic C57BL/6J background. Mice were housed under standard laboratory conditions with 12:12 light/dark cycles and provided with food and water *ad libitum*. All experiments were conducted in accordance with the guidelines from the National Institutes of Health and the University of Wisconsin-Madison Institutional Animal Care and Use Committee.

### Bladder and UGT measurements

Mice were euthanized with carbon dioxide followed by cardiac puncture. Three caliper measurements (h, w, d) were taken for each bladder, and volume was calculated by 4/3·π·((x·y·z)/8). Urogenital tracts (UGT) were microdissected and weighed as previously described[23].

### Void Spot Assays

Void spot assays (VSA) were performed as previously described[22]. Briefly, mice were placed on a 16 x 26- cm Grade 238 chromatography paper (Ahlstrom, Kaukauna, WI) for 4 hours with access to food but not water. Assays were performed during the light phase beginning at 10AM. Filter papers were allowed to dry overnight and then imaged using the UVP ChemStudio AnalytikJena (AnalytikJena, Germany) imager under ultraviolet light using an ethidium bromide filter and 500 ms exposure. Images were imported into ImageJ (v.1.46r) and void spots analyzed using the VoidWhizzard plugin[24]. Data were analyzed by an individual blinded to mouse genotype.

### Uroflowmetry

Uroflowmetry was performed as previously described[25, 26]. Briefly, mice were acclimated to the testing room for 1 hour and then placed in uroflowmetry chambers for 4 hours with access to water but not food. Uroflowmetry data was collected using Raspberry Pi cameras and processors (RS Components Limited, Corby, UK) as previously described[26]. Data was analyzed by an individual blinded to mouse genotype. Urination events in which urine hit the bars of the metabolic chamber floor were excluded from the analysis. A scale from 0 to 3 was used to determine a numerical value for stream rating, in which a rating of 0 corresponded to a single droplet void or significant time between drips and 3 being a void of a strong stream of urine.

### Urethane anesthetized cytometry

Urethane-anesthetized cytometry was performed as previously described[25, 27]. Briefly, mice were anesthetized using a subcutaneous injection of urethane (AC325540500, Thermo Fisher Scientific, Waltham, MA, USA) at a dose of 1.43 g/kg in saline. A PE-50 tubing catheter (NC9140178, Thermo Fisher Scientific, Waltham, MA, USA) was surgically sutured into place at the bladder dome. Saline was infused into the bladder at a rate of 0.8 mL/hr as previously described[25]. Bladder pressure was recorded using a MLT844 physiological transducer (ADInstruments, Colorado Springs, CO, USA) connected to a FE221 Bridge Amp (ADInstruments, Colorado Springs, CO, USA) with a PowerLab 2/26 (PL2602) data acquisition system. Cystometrograms were recorded for approximately 1 h, or until a consistent pattern of voiding was achieved. A total of 5 consecutive voids for each mouse were averaged and analyzed for data collection and selected by an individual blinded to mouse genotype. Measured parameters included void duration, intervoid interval, baseline pressure, normalized threshold pressure, normalized maximum voiding pressure, non-voiding contractions, and compliance.

### Histology

Mice were anesthetized with an excess of isoflurane. Following cessation of respiration, mice were transcardially perfused with PBS followed by 4% paraformaldehyde. Spinal cords and relevant tissues were fixed in 4% paraformaldehyde for 24 hours. Spinal cords were decalcified in 14% EDTA for two weeks, with EDTA changes every 48 hours. For OCT-embedded tissues, tissues were rehydrated in 40% sucrose for 48 hours, embedded in Tissue-Tek OCT Compound, frozen on dry ice, and stored at –80C. Tissues were sectioned on a Leica CM1800 cryostat at section thickness of 60 µm and mounted on Superfrost Plus microscope slides. The slides were stored at -80°C. For paraffin-embedding, tissues were dehydrated in 70% ethanol for 24 hours prior to paraffin-embedding and sectioning; sections were stored at room temperature.

### Immunofluorescence and confocal microscopy

For immunofluorescence, slides were thawed and then washed with PBS with 1% bovine albumin serum (BSA) for 5 minutes. Blocking buffer with 0.1% Triton X was then added for one hour. For slides stained for amyloid-beta, sections were stained with rabbit anti-amyloid-beta (1:100, D54D2, Cell Signaling Technology), Tubulin Beta-3 Chain (TUBB3) (1:250, Biolegend), Podoplanin (1:250, eBio8.1.1, eBioscience), and DAPI. For slides stained for glial fibrillary acidic protein (GFAP), slides were stained with GFAP (1:100, AB5541, Sigma-Aldrich), rabbit anti-amyloid-beta (1:100, D54D2, Cell Signaling Technology), and DAPI. Slides were also stained with cleaved caspase-3 (1:200, Asp175, Cell Signaling Technology), Podoplanin (1:250, eBio8.1.1, eBioscience), and TO-PRO3 (T3605, Invitrogen). Slides were stained with Anti-Vesicular Acetylcholine Transporter (VAchT) (1:500, ABN100, Sigma Aldrich), c-FOS (1:400, 9F6, Cell Signaling Technology), and TO-PRO3 (T3605, Invitrogen). Slides were stained overnight at 4°C. Following staining with primary antibody solutions, sections were washed three times for ten minutes in blocking buffer, and slides were incubated with the appropriate secondary antibody staining solution for two hours at room temperature. Secondary antibodies used were donkey anti-rabbit AlexaFluor568 (1:500, Invitrogen, A10042), donkey anti-goat AlexaFluor488 (A32814, 1:500), goat anti-chicken AlexaFluor488 (A11039, 1:500), and donkey anti-rabbit AlexaFluor647 (1:500, Invitrogen, A31573). Following secondary staining, sections were washed three times for ten minutes and mounted with Prolong Gold mounting medium with DAPI or Prolong Gold without DAPI. Images were acquired using an Olympus Fluoview FV1200 confocal microscope and a Nikon A1R Confocal Microscope. Laser imaging settings were kept the same across the replicates. Images were analyzed using FIJI software utilizing the same brightness and contrast levels.

### Congo red staining and polarized microscopy

Congo red staining was performed on decalcified, whole-head sections embedded in paraffin. Sections were deparaffinized, stained with Harris hematoxylin, rinsed with 95% ethanol, stained with Alkaline Congo Red Solution for 20 minutes, rinsed with NaCl ethanol followed by absolute alcohol twice, cleared with xylene, and mounted in synthetic mounting media. Images were acquired using a Nikon Eclipse E600 microscope fitted with polarized light filters and a Nikon DSFi2 camera. Images were taken at 20X resolution.

### Statistical analysis

Statistical analysis was performed using GraphPad Prism. Data is presented as ± S.E.M. For image-based analyses, multiple images or sections from the same animal were averaged to generate one value per animal. Where indicated, an unpaired Student’s t-test or one-way ANOVA was performed. Statistical significance was defined as p <0.05. No samples or outliers were excluded; outliers were measured using ROUT outlier test on GraphPad Prism.

## Results

### AD mice exhibit lower urinary tract dysfunction compared to WT controls

To assess for voiding dysfunction in transgenic adult male APP/PS1 mice (AD) and their WT controls, void spot assay was performed. Representative VSA images showed modest urination in WT control mice, whereas AD mice displayed a visibly denser pattern of urination characterized by numerous small voids (**Figure 1A**). Quantification of VSA data showed that AD mice had a significantly greater number of total void spots compared to WT mice (**Figure 1B**). AD mice also exhibited an increase in total urine coverage area (**Figure 1C**). Despite increases in total void spots and void area, spatial distribution of spots did not significantly differ between the two groups. AD and WT mice had a similar distribution of urine spots in the center of the VSA paper (**Figure 1D**) and in the corners (**Figure 1E**). Spot-size binning revealed that an increase in total spot count was primarily a result of a significant increase in the number of spots that fell into the smallest bin size (0.01-0.05 cm^2^) in AD mice (**Figure 1F**).

**Figure 1.**
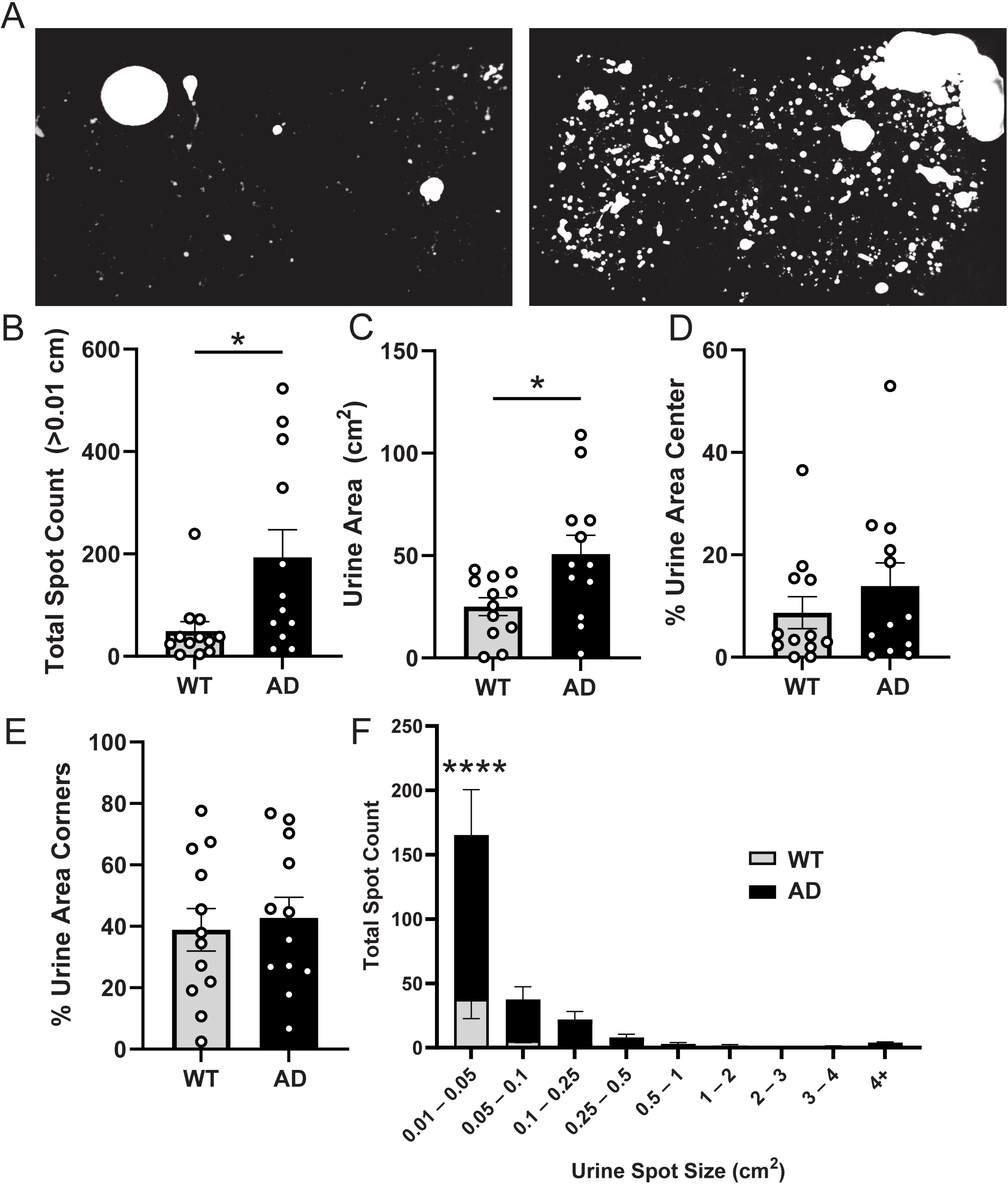
AD mice exhibit increased lower urinary tract dysfunction compared to WT controls. **A)** Representative void spot assay (VSA) images comparing WT controls (left) and transgenic AD (right). **B-E)** Quantification of VSA testing parameters including total spot count **(B)**, urine area **(C)**, percent urine area in center **(D)**, and percent urine area in corners **(E)**. **F)** Distribution of urine spot sizes. Data are mean ± SEM. * p < 0.05, *** p < 0.001, n=9-11.

While the VSA is helpful in assessing overall urinary function over a period of 4 hours, it cannot assess changes in voiding mechanics. We subjected AD and WT control mice to urinary function testing using uroflowmetry, a clinically relevant method that can assess individual void characteristics. Uroflow data showed that AD mice produce smaller and shorter voiding events than WT mice (**Figure 2A-D**). Void mass appeared reduced in AD mice compared to WT controls, although this did not reach statistical significance. Void duration was significantly reduced in AD mice (**Figure 2B**). Quantification showed no significant difference in flow rate (**Figure 2C**) or stream rating (**Figure 2D**).

**Figure 2.**
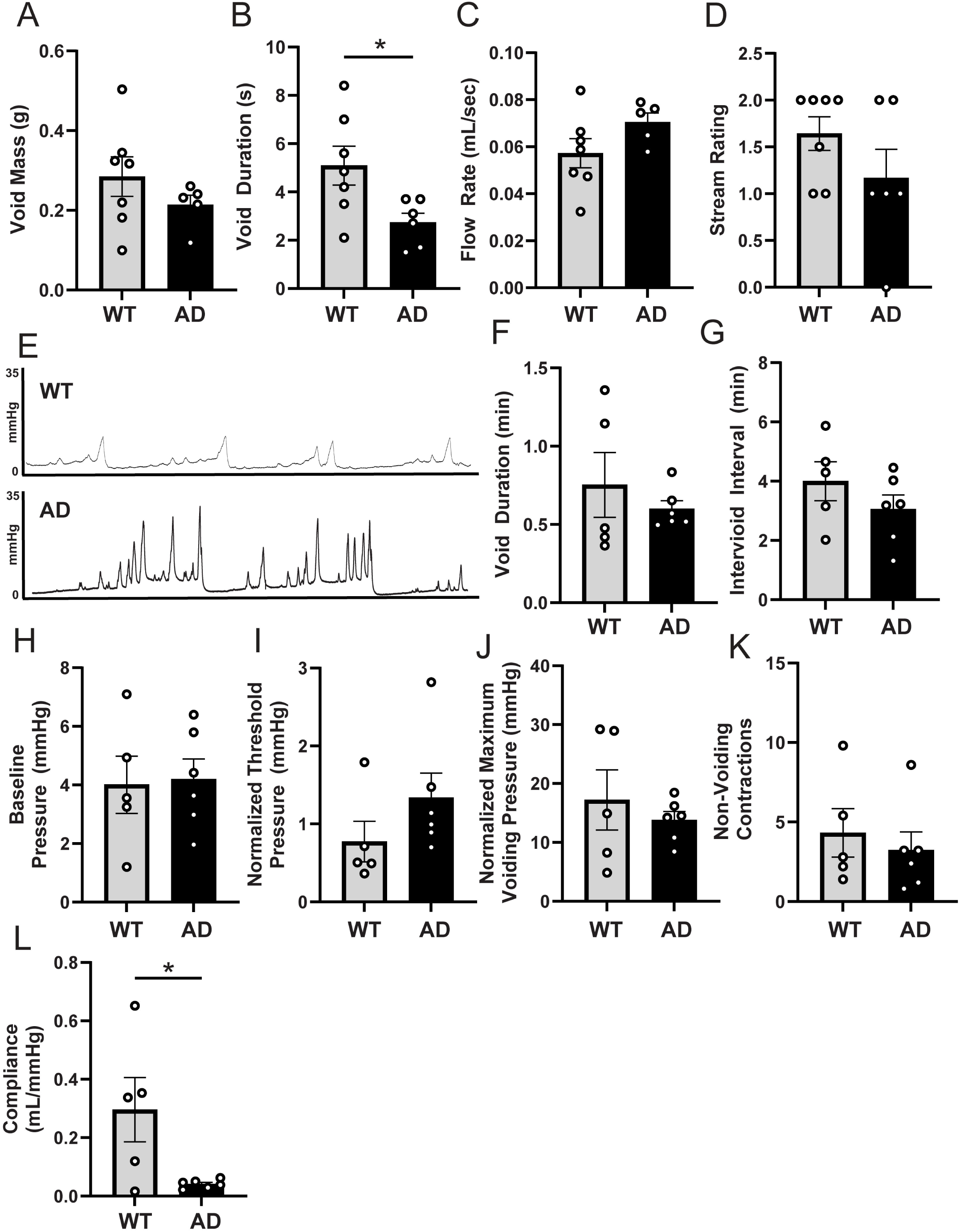
AD mice exhibit altered voiding characteristics on uroflowmetry and bladder dysfunction on urethan-anesthetized cystometry compared to WT controls. A-D) Quantification of uroflowmetry parameters including void mass **(A)**, void duration **(B)**, flow rate **(C)**, and stream rating **(D)**. **E)** Representative cystometrograms from WT control (top) and AD (bottom). F-L) Quantification of cystometrogram parameters including void duration **(F)**, intervoid interval **(G)**, baseline pressure **(H)**, normalized threshold pressure **(I)**, normalized maximum voiding pressure **(J)**, non-voiding contractions **(K)**, and detrusor compliance **(L)**. Data are mean ± SEM. * p < 0.05, ** p<0.01, n=5-6.

To assess bladder storage and voiding mechanics, we performed urethane-anesthetized cystometry (CMG) on AD and WT control mice. Analysis of CMG showed no significant differences in void duration, intervoid interval, baseline pressure, normalized threshold pressure, normalized maximum voiding pressure, or non-voiding contractions (**Figure 2E-K**). Interestingly, in contrast, bladder compliance was significantly reduced in AD mice compared to WT controls (**Figure 2L**).

Taken together, these data assessing urinary health and voiding function suggest that aged adult male AD mice exhibit urinary dysfunction, characterized predominantly by urinary spotting or dribbling, shortened void durations with an overall preserved flow rate, and significantly reduced bladder compliance. These functional changes were not accompanied by significant differences in bladder mass, bladder volume, bladder volume/body mass ratio, or prostate lobe enlargement (**Supplementary Figure 1**).

### AD mice have amyloid-beta deposition at each level of the spinal cord

To investigate the contribution of lower-motor neuron involvement in urinary dysfunction, the spinal cords of aged adult AD mice and their WT controls were collected and decalcified. To define amyloid-beta localization within the spinal cord, the neuronal marker of TUBB3 was used. To test whether amyloid-beta presence is due to lymphatic clearance, podoplanin was examined as it is a widely used marker of lymphatic vessels. Amyloid-beta was found at each level of the spinal cord – cervical, thoracic, lumbar, and sacral (**Figure 3B-E**). Amyloid-beta was detected within the spinal cord parenchyma, particularly in the ventral and dorsal horns, as well as within the dorsal root ganglia, ventral roots, and leptomeningeal/vascular structures (**Figure 3F-I**). Amyloid-beta was observed in association with the anterior spinal artery and other vascular structures, consistent with a vascular distribution that may reflect amyloid angiopathy-like pathology, and was also detected near the central canal (**Figure 3H**). The spinal cord terminates around L1/L2 in mice, which is an area known as the conus medullaris. After the conus medullaris, the separate spinal nerves create the cauda equina. Within the cauda equina, amyloid-beta was found within the spinal nerves root as well at the area of the filum terminale (**Figure 3I**), which is a fibrous filament extending from the conus medullaris to the coccyx to anchor the spinal cord.

**Figure 3.**
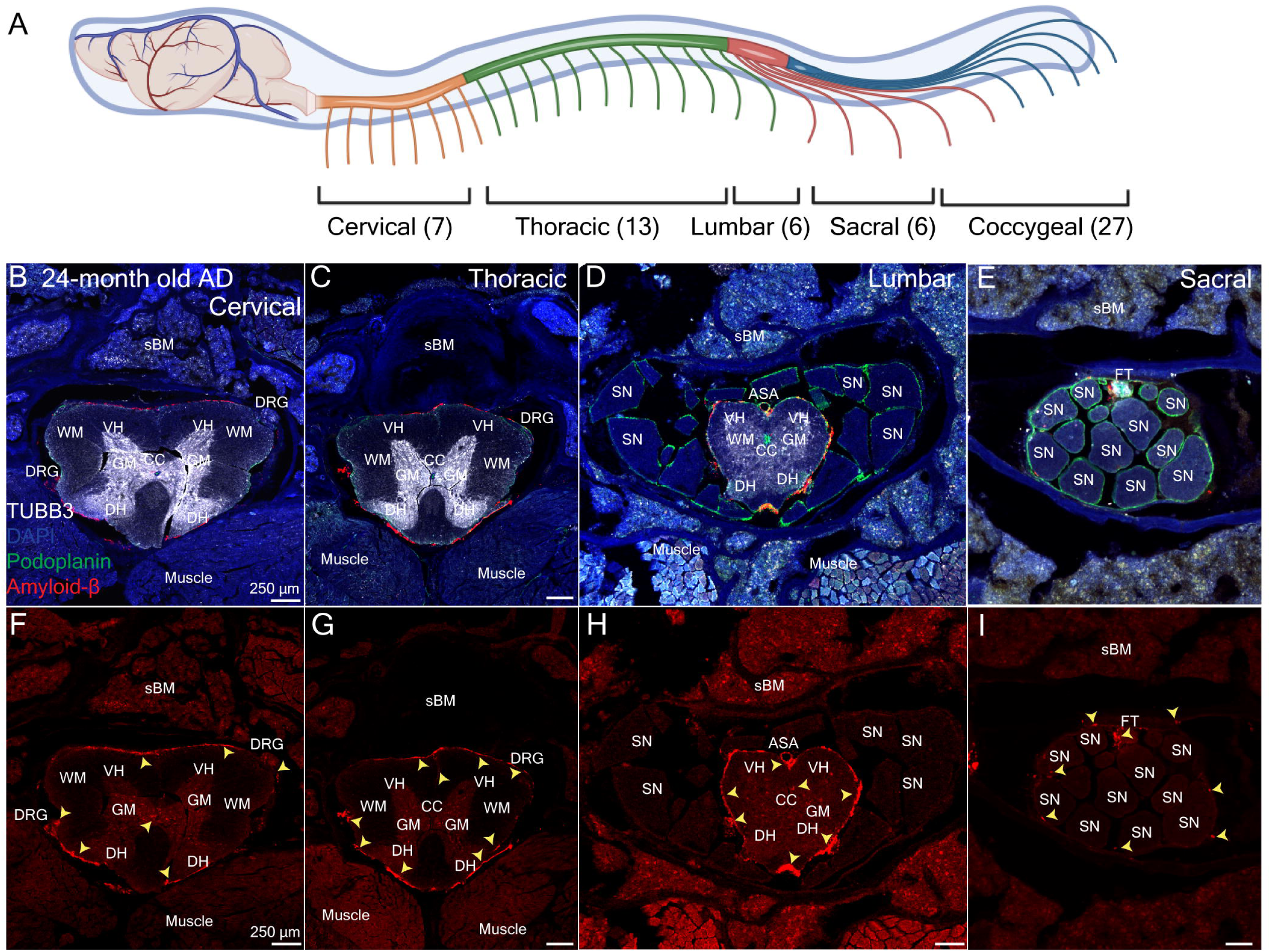
Amyloid-beta is present within the spinal cord in aged AD mice. **(A)** Schematic of mouse vertebral column and corresponding spinal cord regions illustrating the number of vertebrae in each area. **(B-E)** Representative immunofluorescence images of cervical **(B)**, thoracic **(C)**, lumbar **(D)**, and sacral **(E)** spinal cord segments of 24-month-old APP/PS1 stained with TUBB3 (TUJ1), Podoplanin (eBioscience8.1.1), Amyloid-beta (D52D4), and DAPI. **(F-I)** Single channel Amyloid-beta (D52D4) images in the cervical **(F)**, thoracic **(G)**, lumbar **(H)**, and sacral **(I)** regions of the spinal cord. Images are representative of n=4 mice. sBM = spinal cord bone marrow, GM = gray matter, WM = white matter, DH = dorsal horn, VH = ventral horn, DRG = dorsal root ganglion, ASA = anterior spinal artery, SN = spinal nerve, CC = central canal, and FT = filum terminale.

### AD mice have amyloid-beta deposition in areas innervating the bladder

Cerebrospinal fluid (CSF) is present within the central cord of the canal, the subarachnoid space surrounding the spinal cord, and in perineural routes along spinal nerves as they exit the subarachnoid space. The central canal of the spinal cord is lined with ciliated ependymal cells driving CSF movement. It is established that amyloid-beta is present in the subarachnoid space and in plasma, which has provided the basis for a robust field of CSF and serum biomarkers. We hypothesized that amyloid-beta would be present in areas in contact with cerebrospinal fluid such as the central canal, subarachnoid space surrounding the spinal cord, and the perineural space as spinal nerves exit the spinal cord where the fluid is ultimately collected in lymphatic vessels.

The spinal cords of aged adult AD mice and their WT controls were collected, decalcified, and stained for the neuronal marker of TUBB3, amyloid-beta, and podoplanin. Amyloid-beta was found within the central canal of AD mice (**Figure 4B**). To confirm the presence of amyloid-beta within the central canal, the spinal cord of aged adult AD mice and their WT controls were collected, decalcified, and paraffin embedded. The central canal is shown on hematoxylin and eosin staining in AD mice (**Figure 4C**) and WT controls (**Figure 4E**). Adjacent sections were stained with Congo Red, which stains beta-pleated sheets that form the structure of amyloid-beta and will display an “apple-green” birefringence under polarized light. Within the central canal, AD mice had apple-green birefringence displaying amyloid-beta within the central canal (**Figure 4D**) and the WT control did not have any observable staining (**Figure 4F**). Next, the filum terminale at the level of the coccyx was examined. Amyloid-beta was found in the area of the filum terminale (**Figure 4G-H**) and was not present in the control tissue (**Figure 4I**).

**Figure 4.**
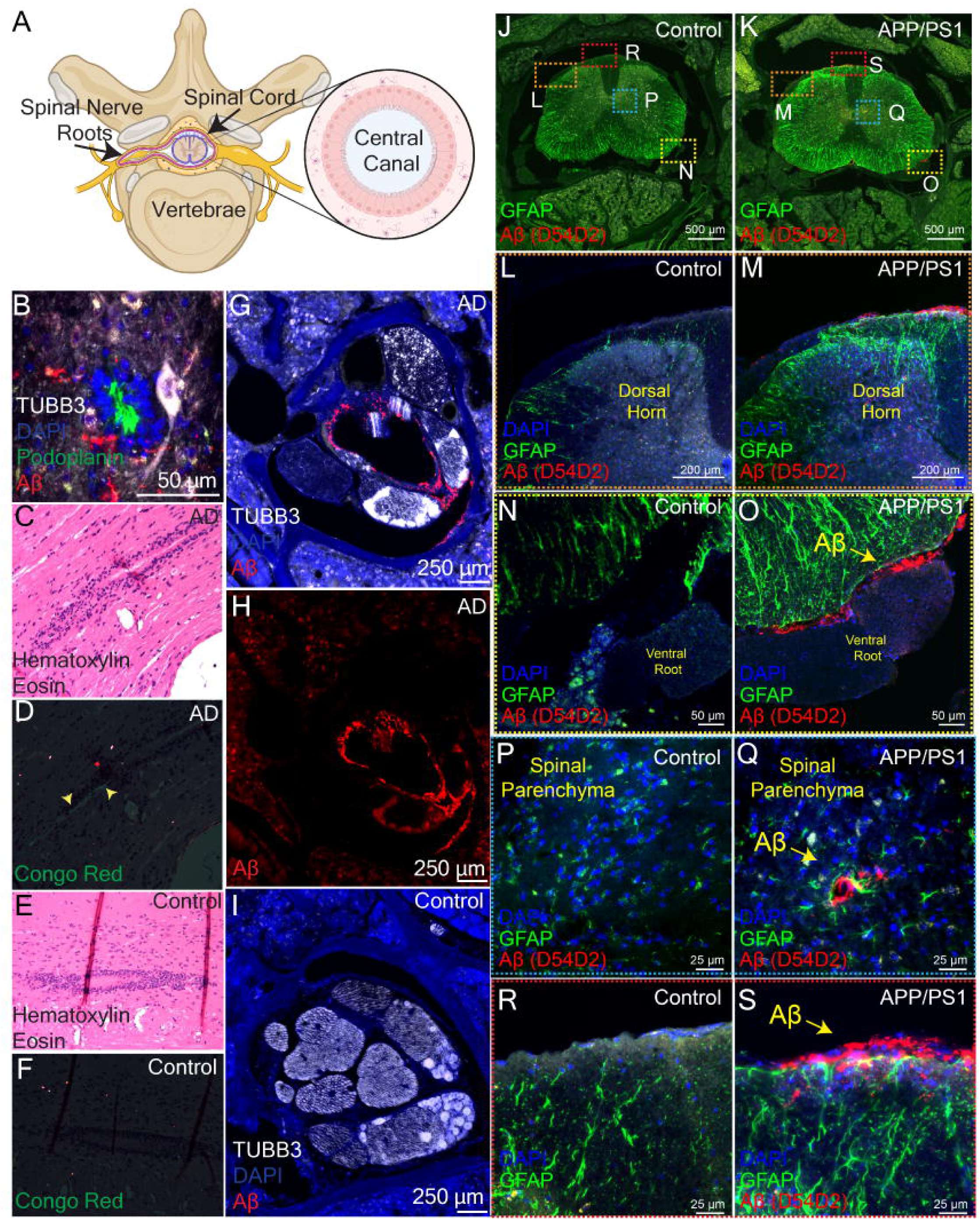
Amyloid-beta within the spinal cord localizes to areas of cerebrospinal fluid efflux and neuronal signaling. **(A)** Anatomical schematic illustrating transverse view of the spinal cord displaying cerebrospinal fluid (CSF) present in the central canal, spinal cord, dorsal root ganglion, and ventral root ganglion. **(B)** Representative magnified immunofluorescence image of the central cord stained with TUBB3 (TUJ1), Podoplanin (eBioscience8.1.1), Amyloid-beta (D52D4), and DAPI of a 24-month-old APP/PS1 mouse. Representative image of n=4. **(C-D)** Representative images from 24-month-old APP/PS1 **(C-D)** and 24-month-old littermate control **(E-F)** with images from slides stained with hematoxylin and eosin **(C and E)** to display histology and images taken under polarized light **(D and F)** displaying apple green birefringence **(D). (G-I)** Representative immunofluorescence image of the filum terminale stained with TUBB3 (TUJ1), Amyloid-beta (D52D4), and DAPI. **(J-S)** Representative immunofluorescence images of 24-month-old APP/PS1 **(K, M, O, Q, S)** and 24-month-old littermate control **(J, L, N, P, R)** stained with TUBB3 (TUJ1), Amyloid-beta (D52D4), glial fibrillary acidic protein (GFAP), and DAPI displaying localization in the dorsal horn and dorsal root ganglion area **(L-M)**, ventral root ganglion **(N-O)**, spinal parenchyma **(P-Q)**, and spinal meningeal area **(R-S)**. Images are representative of n=4.

Next, we examined the presence of amyloid-beta in areas important for neuronal signaling to the bladder. The spinal cords of aged AD and their WT controls were collected, decalcified, and stained for glial fibrillary acidic protein (GFAP), amyloid-beta, and DAPI. Amyloid-beta was present in several areas responsible for bladder signaling including the dorsal horn (**Figure 4M**), ventral root (**Figure 4O**), and the spinal parenchyma (**Figure 4Q**). Amyloid-beta was also consistently seen in the meningeal layer of the spinal cord (**Figure 4S**). Importantly, control tissue did not show any staining as expected (**Figure 4J, 4L, 4P, and 4R**). Greater GFAP staining appeared in areas adjacent to amyloid-beta deposition, but this trend was not significant (**Figure 4J-S**).

### AD mice have decreased neuronal transmission and activation in areas innervating the bladder

It has been established that amyloid-beta is a neurotoxic protein[28–31]. As we found amyloid beta present in many areas important for neuronal signaling, we investigated if there were impacts on neuronal transmission and activation. Parasympathetic innervation to the bladder governs micturition with sacral preganglionic neuronal cell bodies sending acetylcholine as a neural transmitter through the ventral nerve root and the cauda equina to reach the postganglionic neurons on the bladder to signal detrusor contraction and micturition. To identify cholinergic innervation around motor neurons and potential changes in cholinergic circuitry, VAchT was stained together with the far-red fluorescent nuclear counterstain TO-PRO3. Furthermore, to test if amyloid-beta deposition induced changes in neuronal signaling, the early cellular activation marker, c-Fos, was examined (**Figure 5A-C**). Aged AD mice had significantly reduced VAchT immunoreactivity within the cauda equina in comparison to age-matched controls (**Figure 5D**). In addition, aged AD mice had significantly reduced c-FOS in this region (**Figure 5E**). To understand if the nerves had undergone apoptosis, decalcified spinal cords were stained for cleaved caspase-3, podoplanin, and TO-PRO-3, a nuclear marker (**Figure 5G-I**). Expression of cleaved caspase-3 was found within the gray matter (**Figure 5G-H**) and around the anterior spinal artery (**Figure 5I**). Surprisingly, the aged WT control had expression of cleaved caspase-3 in the gray matter (**Figure 5G**). Apoptosis within the gray matter generally occurs due to trauma or injury; recent research has suggested that with age there is an increase of cleaved caspase-3 in the gray matter[32]. In comparison of the AD mice to WT mice, there was a trend of increased cleaved caspase-3 (**Figure 5F**) but it was not significant.

**Figure 5.**
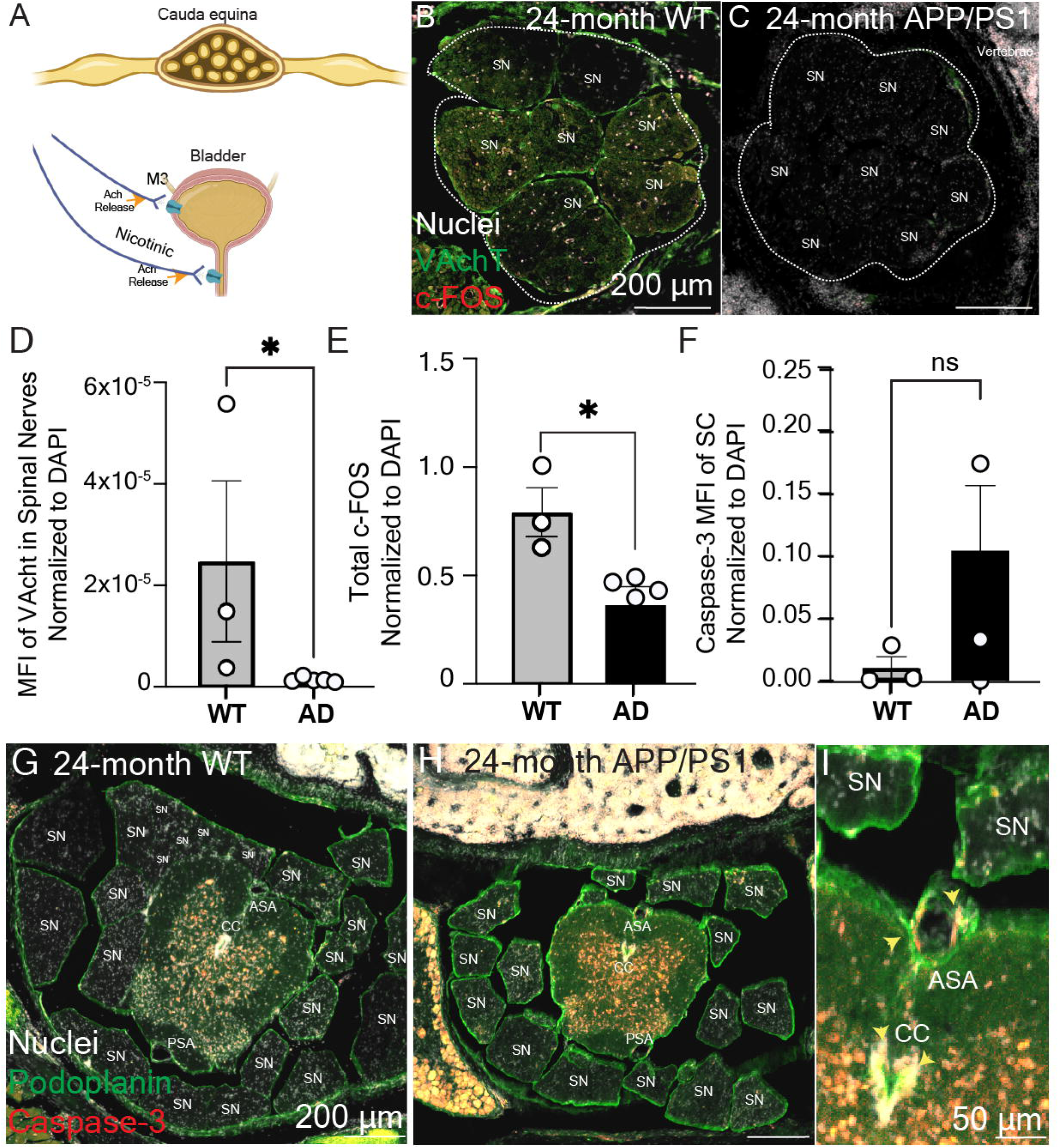
AD mice have decreased neuronal activation and acetylcholine release in the lumbosacral region responsible for bladder innervation. **(A)** Anatomical schematic of bladder innervation displaying sympathetic innervation pathways originating from spinal nerve roots in the cauda equina resulting in acetylcholine release at muscarinic receptors on the bladder smooth muscle and acetylcholine release at nicotinic receptors on the detrusor. **(B-C)** Representative immunofluorescence images of 24-month-old littermate **(B)** and APP/PS1 mice **(C)** stained with vesicular acetylcholine transporter (VAchT), c-FOS, and TO-PRO3, a nuclear marker. Representative image of n=3-5 mice. **(D-F)** Quantification of mean fluorescence intensity (MFI) of VAchT normalized to DAPI **(D)**, c-FOS normalized to DAPI **(E)**, and cleaved caspase-3 **(F)**. Results of an unpaired student’s t-test. **(G-H)** Representative immunofluorescence images of 24-month-old littermate **(G)** and APP/PS1 mice **(H)** stained with cleaved caspase-3 (Asp175), Podoplanin, and TO-PRO3, a nuclear marker). **(I)** Magnified images showing caspase-3 signal near central cord and anterior spinal artery. Representative image of n=3-5. SN = spinal nerve, CC = central canal, ASA = anterior spinal artery, PSA = posterior spinal artery.

## Discussion

Alzheimer’s disease (AD) is expected to increase in prevalence from 65 million individuals worldwide to 82 million by 2050 and already represents a growing public health crisis with large psychosocial and economic costs[1]. Urinary incontinence (UI) is a common comorbidity in AD and has significant impacts on quality of life for patients and their caregivers[11]. While UI in dementia has historically been hypothesized to result from functional incontinence, urodynamic studies in dementia populations support that urinary dysfunction may be as result of disease-mediated pathology rather than purely functional in nature[33]. Moreover, the clinical management of UI in AD represents a unique therapeutic paradox, as pharmacologic strategies for dementia and UI target acetylcholine signaling in directly opposing directions. Acetylcholinesterase inhibitors used to slow cognitive decline dementia can lead to symptoms of overactive bladder and lower urinary tract dysfunction, while antimuscarinics used to treat UI have been shown to increase risk of dementia and worsen cognitive decline in older individuals[14, 34]. Collectively, these observations support that urinary dysfunction in AD is likely multifactorial and can include urge incontinence and mixed phenotypes in addition to functional incontinence. However, the underlying pathophysiology driving urinary dysfunction in AD remains largely unknown. To investigate this further, we present a combination of functional voiding assays in aged AD mice, detailed characterization of amyloid-beta deposition patterns in mouse spinal cord and peripheral structures relevant to bladder control, and analysis of neuronal signaling markers in bladder-innervating pathways.

Neuronal control micturition is a highly organized and complex system that integrates input from supraspinal micturition centers, the spinal cord, and peripheral afferent and efferent pathways[35]. Disruption at distinct levels produces characteristic lower urinary tract phenotypes[36]. Suprasacral neurologic disruption, for example from spinal cord injuries or multiple sclerosis lesions, commonly produces storage-phase abnormalities including detrusor overactivity, whereas sacral spinal cord, nerve root, or peripheral nerve involvement can compromise parasympathetic signaling and impair voiding function, often manifesting as underactive bladder or overflow incontinence phenotype[37–39]. Importantly, incomplete or partial disruption in these pathways can generate complex phenotypes that do not readily map onto a single diagnosis or incontinence type, highlighting the need for multimodal functional testing in preclinical models[39].

Urodynamic assessment of urinary function in aged AD mice revealed a pattern of lower urinary tract dysfunction characterized by increased urinary frequency, an increased number of small-volume voids or urinary dribbling, shortened voiding duration, and significantly reduced bladder compliance. One limitation of the study is that water consumption was not measured in mice, thus it is possible that the degree of water consumption contributed to the observed urinary phenotype. However, population studies in AD patients have shown that patients have increased dehydration rather than increased water consumption[40, 41]. Importantly, flow rate, stream rating, and voiding pressures were largely unchanged compared to their WT controls, and these functional changes occurred in the absence of bladder enlargement, increased bladder mass, or enlargement of any of the prostate lobes. Together, these findings argue against a primary bladder outlet obstruction and instead suggest a neurologic contribution to urinary dysfunction. The findings of small-volume voiding and reduced compliance are most consistent with a storage-phase abnormality, while shortened void duration may reflect impaired coordination or early termination of voiding rather than classic detrusor underactivity[42–44].

Although some features of the urinary phenotype in the aged AD mice resemble overactive bladder, the overall pattern does not fully align with classic detrusor overactivity. Specifically, cystometric analysis did not demonstrate increased non-voiding contractions or elevated baseline or threshold voiding pressures. A trend toward reduced voided volume and significantly shortened voiding duration suggest possible impaired detrusor contraction or altered detrusor external sphincter coordination[45]. However, in our study preserved flow rate with no significant increase in voiding pressures argues against detrusor external sphincter dyssynergia as a primary driver of the phenotype. Although we cannot exclude subtle sphincter discoordination without formal analysis of external urethral sphincter function.

Mixed features such as these can occur in neurogenic bladder phenotypes involving partial disruption of spinal or peripheral pathways and are described in conditions affecting the cauda equina or sacral nerve roots[46–48]. The absence of post-void residual measurements represents a limitation of this study, as residual volume would be required to determine whether detrusor overactivity with impaired contractility (DOIC) contributes to the observed phenotype[49]. Interestingly, retrospective clinical data indicate that mixed storage-voiding phenotypes, including DOIC, are present in a subset of patients with AD[50, 51].

Given the complex neuronal control of micturition and the observed functional phenotype, we examined the thoracolumbar and lumbosacral spinal cord as a potential site of dysregulation. Amyloid-beta deposition was consistently identified throughout the spinal cord in aged AD mice, including cervical, thoracic, lumbar, and sacral segments, and within anatomical structures critical for bladder innervation such as the dorsal root ganglia, ventral roots, conus medullaris, and cauda equina. These deposits were found within the spinal parenchyma and meningeal layers surrounding these structures. The presence of amyloid-beta within the central canal, filum terminale, and meningeal layers, regions that are in direct contact with cerebrospinal fluid (CSF), raises the possibility that spinal amyloid deposition may occur through CSF-mediated mechanisms, although how these deposits are transported and accumulate in these areas are unknown. Studies have shown that amyloid-beta deposits are neurotoxic to ciliated ependymal cells, leading to ciliary dyskinesia, decreased CSF clearance, and disruption of the blood-CSF barrier[52–54]. An important consideration for the interpretation of these findings is that the amyloid-beta antibody used detects amyloid-associated species, but it does not distinguish between soluble and insoluble amyloid-beta or between diffuse and fibrillar deposits. There is active research that has indicated that certain forms of amyloid-beta may have different consequences such as soluble amyloid-beta being more neurotoxic while insoluble amyloid-beta is more prone for aggregation and plaque formation[28, 30, 31, 55, 56]. Therefore, while amyloid immunoreactivity was observed throughout the spinal cord and bladder-associated neuronal pathways, the precise nature of these deposits remains to be determined. Future studies defining the biochemical and structural properties of these deposits may provide insight into how specific amyloid species contribute to spinal cord dysfunction and urinary symptoms in AD.

In these studies, we demonstrate amyloid-beta deposition at the meningeal layers associated with prominent Podoplanin expression indicating the involvement of podoplanin-positive lymphatic vessels in amyloid-beta transport (Fig 3D). Additionally, occasional Podoplanin positive perineural spaces contained amyloid-beta (Fig 3E), suggesting the role of perineural drainage pathways for amyloid-beta distribution. Dissecting the amyloid-beta drainage pathways throughout the spinal cord using further lymphoid vessel specific markers will shed light for sources of amyloid-beta in the spinal cord and designing targeted therapies.

To assess whether amyloid-beta deposition was associated with altered neural function, we evaluated markers of neuronal signaling and activation in bladder-innervating spinal pathways. Aged AD mice exhibited reduced vesicular acetylcholine transporter (VAchT) expression within the cauda equina. Concurrently, data showed a significant reduction in total c-Fos expression. Together these findings suggest alterations in neural pathways associated with bladder control in aged AD mice. As the cauda equina and spinal nerves predominantly contain axons and associated glial cells; therefore, additional investigation will be required to establish how changes in VAChT and c-Fos immunoreactivity relate to bladder-innervation and function. Nevertheless, these findings suggest altered activity-dependent cellular responses within bladder-associated neural pathways. Further studies examining neuronal cell bodies and distal bladder innervation will help define the cellular origin and functional relevance of these changes to bladder innervation. However, these findings did not suggest a total decrease in neuronal mass, as caspase-3, a marker of apoptosis, was not significantly increased compared to WT controls. Therefore, we hypothesize that there may be a functional impairment or reduced recruitment of micturition-related nerve pathways in the spinal cord of aged AD mice compared to their WT controls. Such changes are consistent with neurogenic bladder conditions in which partial disruption of afferent or efferent signaling leads to inefficient bladder emptying and altered storage function.

Collectively, these data suggest that urinary dysfunction in aged AD mice may correlate with the dysregulation within spinal and peripheral components of the micturition pathway rather than purely functional incontinence, which is incontinence driven by other underlaying conditions. Importantly, our findings do not establish that spinal amyloid-beta deposition is the primary driver of urinary dysfunction but instead identify a novel anatomical and functional association that warrants further mechanistic investigation. Future studies incorporating longitudinal assessment, direct measures of sensory afferent signaling, post-void residual volume, and targeted manipulation of spinal cord pathways will be necessary to define causality and to determine to what extent extracerebral amyloid-beta deposits contribute to urinary dysfunction in AD.

Future directions include increasing our analysis to examine post-mortem human samples. Few studies have reported amyloid-associated neuropathology in the spinal cord of human AD patients, most being limited to leptomeningeal, meningeal, or vascular deposits associated with cerebral amyloid angiopathy [3, 57, 58]. However, there have been several studies reports on hyperphosphorylated tau within the spinal cord parenchyma with implications for chronic pain and impaired tactile sensation[59–62]. Interestingly, in a study of post-mortem spinal cords, the presence of hyperphosphorylated tau was most abundant within the cervical spinal cord followed by the thoracic, lumbar, and sacral[63]. Although not in the context of AD, other work has shown evidence of Aβ42 deposition in the lumbar spinal cord, specifically within the anterior horn, of patients with amyotrophic lateral sclerosis [64]. It will be important to see if the amyloid-beta deposition in aged APP/PS1 mice that we report here correlate with AD patient results.

In summary, this study demonstrates that aged AD mice develop a pattern of lower urinary tract dysfunction characterized by urinary frequency, shortened void duration, and reduced bladder compliance in the absence of mechanical bladder outlet obstruction. Amyloid-beta deposition and altered neuronal signaling were identified within spinal regions involved in bladder and external urethral sphincter control in aged APP/PS1 mice. These findings help to explain our current understanding of urinary dysfunction in AD and suggest that spinal and peripheral nerve pathways should be considered alongside cortical and functional mechanisms when evaluating and treating urinary symptoms in patients with Alzheimer’s dementia.

## Acknowledgements

The authors would like to thank Dr. Melinda Herbath, Jenna Port, and Dr. Thiunuwan Priyathilaka Thanthrige, Emily Ricke, and Dr. Johanna Hannan, Abigail Sehmer, Avery Holtzman, and Simran Sharma for their helpful discussions and constructive critiques on this project.

## Funding

The authors would like to acknowledge the Cancer Center Support Grant: NCI P30 CA014520, University of Wisconsin Small Animal Imaging & Radiotherapy Facility and NIH S10OD028670-01 for supporting this work. Additionally, we thank the University of Wisconsin Translational Research Initiatives in Pathology (TRIP) Laboratory, supported by UW Department of Pathology and Laboratory Medicine, UWCCC (P30CA14520) and the Office of the Director NIH (S10 OD023526). We thank the Optical Imaging Core, supported by 1S10034394-01. The work was supported by NIH grant(s) 5T32GM140935 to S.M.V; F30DK143709 and T32GM141013 to A.R.L; 5R01NS126595 to Z.F; P30AG062715 to J.J.N; VA GRECC to J.J.N and M.P; and R01DK127081, R01DK131175, and U54DK104310 to W.A.R. The authors have no conflict of interest to declare.

**Supplemental Figure 1. AD mice do not have altered prostate or bladder mass compared to WT controls.** Quantification of body mass **(A)**, bladder volume **(B)**, bladder volume/body mass ratio **(C)**, anterior prostate mass **(D)**, ventral prostate mass **(E)**, and dorsolateral prostate mass **(F)**.

