## Supplementary figures and images for "Amyloid-beta is present in the spinal cord of APP/PS1 mice and may contribute to neuropathology manifesting as lower urinary tract dysfunction"

### Supplemental Figure 1

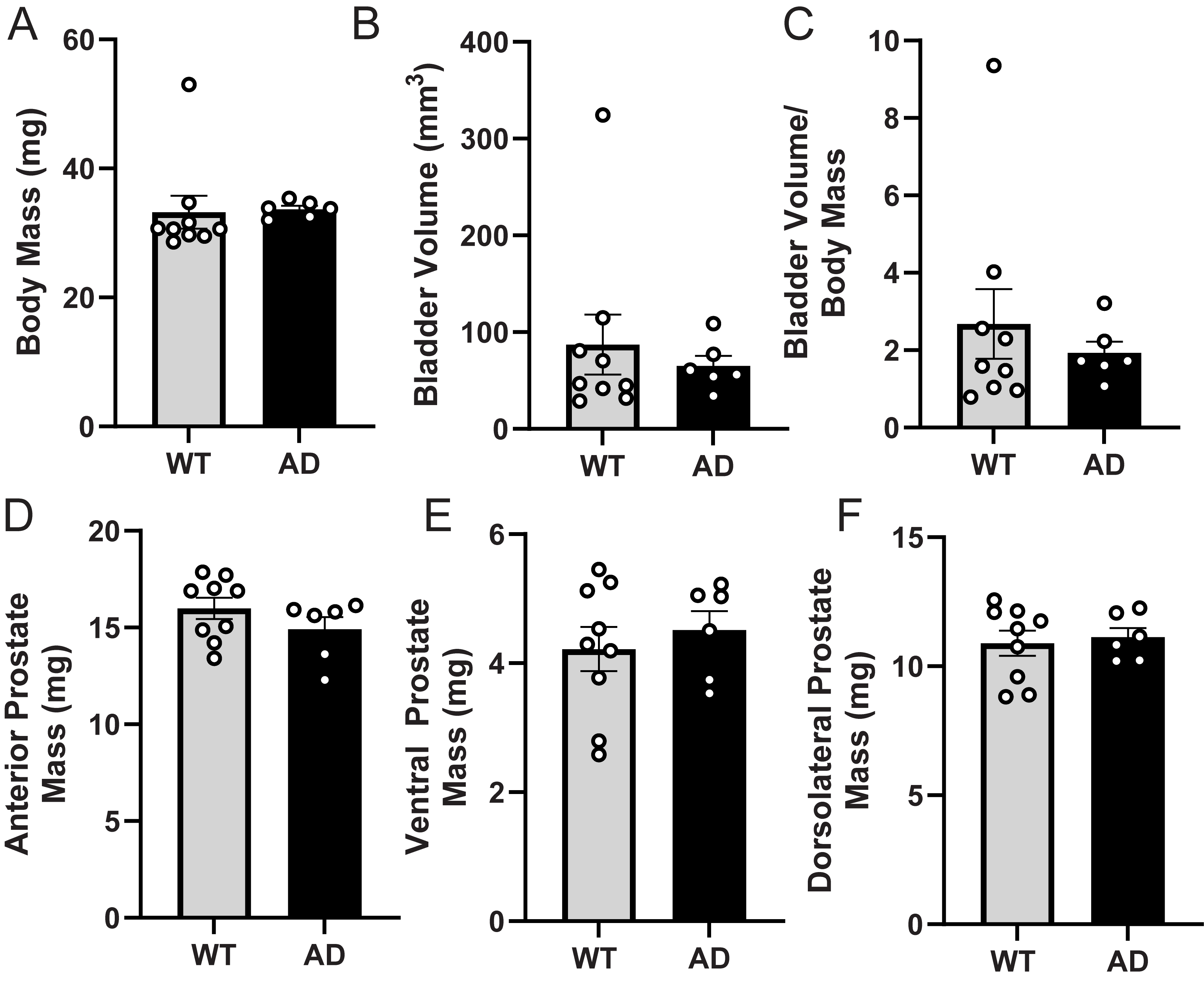
